# Expression patterns of m^6^A RNA methylation regulators under apoptotic conditions in various human cancer cell lines

**DOI:** 10.1101/2023.08.10.551971

**Authors:** Azime Akçaöz-Alasar, Buket Sağlam, Ipek Erdogan Vatansever, Bünyamin Akgül

**Affiliations:** Noncoding RNA Laboratory, Department of Molecular Biology and Genetics, İzmir Institute of Technology, Gülbahçeköyü, 35430 Urla İzmir, Turkey

## Abstract

The emerging evidence suggests that epitranscriptomics changes play a crucial role in the pathogenesis of cancer. However, expression patterns of m^6^A RNA modifiers under apoptotic conditions are unknown. We measured the transcript abundance of m^6^A RNA modifiers under cisplatin- and tumor necrosis factor alpha (TNF-α)-induced apoptotic conditions. In general, the abundance of m^6^A modifiers is increased upon cisplatin treatment whereas TNF-α treatment has led to a reduction in their expression. Specifically, cisplatin-induced apoptosis, but not TNF-α mediated apoptosis, lowered the abundance of METTL14 and FTO transcripts. Additionally, cisplatin treatment plummeted the abundance of IGF2BP2 and IGF2BP3 readers following cisplatin treatment. These results suggest that differential response of cancer cells to apoptotic inducers could be partially due to m^6^A RNA modifiers.

## Introduction

Global cancer incidence in women, as reported by the world health organization, indicates that the most widespread cancer types are breast, colorectal, lung and cervix cancers with a frequency of 25.8, 9.9, 8.8 and 6.9%, respectively (Norpo’latovna, 2023; Siegel et al., 2023). Thus, the ability to identify the prognostic risk factors of cancer types and their growth, progression and treatments have been widely valued by researchers. Although genetic, genomics and biochemical approaches have uncovered numerous molecular mechanisms that contribute to tumorigenesis, recent developments have shown that *N*^6^-methyladenosine (m^6^A) RNA modification is closely associated with activation and inhibition of tumorigenesis (Du et al., 2022; Jiang et al., 2023; Ma et al., 2022; C. Zhang et al., 2022). m^6^A RNA modification refers to the attachment of a methyl group to the 6-carbon of adenosine, which is a co-transcriptional and dynamic regulatory mark catalyzed by a series of enzymes (Leonetti et al., 2020). Writer, erasers, and readers are responsible for attachment, removal, and recognition of m^6^A sites, respectively, modulating a wide range of RNA fates such as splicing, translation, microRNA processing, nuclear export, RNA stability and decay (Akçaöz & Akgül, 2022; Niu et al., 2013). Consequently, m^6^A RNA methylation plays a pivotal role in orchestrating various biological processes such as gene regulation, DNA damage response, signal transduction and apoptosis (Akçaöz-Alasar et al., 2023; Alasar et al., 2022; Z. X. Liu et al., 2018; Roundtree et al., 2017).

The existing evidence clearly suggests that some m^6^A RNA writers, erasers or readers are associated with cancer formation or development (Deng et al., 2022; L. Liu et al., 2022; N. Zhang et al., 2021). Most studies are centered around Fat mass and obesity-associated protein (FTO) and Methyltransferase like 3 (METTL3). FTO acts as an oncogene in acute myeloid leukemia as it increases malignant transformation and tumor formation (Z. Li et al., 2017). Similarly, while FTO mRNA and protein levels are upregulated in human lung cancer tissues, its downregulation reduces cell proliferation and tumor growth (J. Li et al., 2019). The decrease in METTL3 level, on the other hand, impairs tumor growth due to decreased m^6^A methylation and inhibits malignant transformation of glioblastoma (Visvanathan et al., 2018). However, proteins in the m^6^A machinery do not always act as oncogenes, and in some cases, they might be a positive regulator of tumor growth. For example, in gastric cancer with METTL3 overexpression, METTL3 knockdown impairs the cell proliferation and migration capacity (T. Liu et al., 2020). Additionally, METTL3 induces tumorigenesis and growth in hepatocellular carcinoma (Chen et al., 2018). These results clearly suggest that the contribution of m^6^A machinery to cancerogenesis differs depending on the type of cancer. Consequently, a total account of writer, eraser and reader expression in this fundamental process is important to decipher the complex association between m^6^A RNA methylation and cancer. In this study, we measured the abundance of transcripts that orchestrate m^6^A methylation in healthy and different cancer cell lines along with their expression under apoptotic conditions.

## Material & Methods

### Cell Culture

HeLa cells were purchased from DSMZ GmbH (Germany). ME-180 (HTB-33), MDA-MB-231 (HTB-26) and MCF-7 (HTB-22) cell lines were purchased from ATCC (United States). A549, Caco-2, H1299 and HCT116 cell lines were kindly provided by Dr. Serdar Özçelik of Izmir Institute of Technology (Turkey), Dr. Sreeparna Banerjee of Middle East Technical University (Turkey), Dr. Hakan Akça of Pamukkale University (Turkey) and Dr. Semra Koçtürk of Dokuz Eylül University (Turkey), respectively. Caco-2 cells were cultured in EMEM (Gibco, United States) supplemented with 20% FBS, 2 mM L-glutamine, 1 mM Na-pyruvate (Gibco, United States) and 1X non-essential amino acids (Gibco, United States). The other cell lines were maintained in the following conditions: RPMI 1640 (Gibco, United States) for HeLa and H1299 cells, DMEM (Lonza, Switzerland) with high glucose for A549 and MDA-MB-231, McCoy’s 5A (Lonza, Switzerland) for ME-180 and HCT116 cells. Except for EMEM, all media were supplemented with 10% fetal bovine serum (FBS) (Gibco, United States) and 2 mM L-glutamine. HeLa, MCF7 and A549 cell lines were maintained in a humidified atmosphere of 5% CO_2_ at 37°C. 80 µM, 100 µM and 80 µM of cisplatin (Santa Cruz Biotechnology, United States) for 16h and 75 ng/ml TNF-α in 5 μg/ml CHX (Applichem, Germany), 10 ng/ml TNF-α in 5 μg/ml CHX and 20 ng/ml TNF-α in 10 μg/ml CHX of TNF-α (Biolegend, United States) for 24h treatments were performed in varying concentrations, respectively. 0.1% (v/v) DMSO and cycloheximide (CHX) were used as negative controls for cisplatin and TNF-α treatments, respectively. Treated cells were harvested by 1X Trypsin-EDTA (Gibco, United States) and washed in 1X cold PBS (Gibco, United States) prior to staining with Annexin V-PE (Becton Dickinson, United States) and 7AAD (Becton Dickinson, United States) in the presence of 1X Annexin binding buffer (Becton Dickinson, United States) for flow cytometry analysis (Yaylak et al., 2019). Apoptotic and live populations were determined by FACSCanto flow cytometer (Becton Dickinson, United States).

### RNA isolation

Cells were dissolved in an appropriate amount (750 µl for 5×10^6^ cells) of TRIzol^TM^ reagent (Invitrogen, ThermoFischer, Waltham, MA, USA) and stored at −80 [C until isolation. The total RNA isolation was performed following the manufacturer’s procedure. If necessary, 1 µl (20 mg/ul) glycogen was added prior to precipitation of total RNAs at +4 [C and 12,000 xg for 10 minutes. RNA pellets were air-dried for 5-10 minutes and dissolved in 20-30 µl of nuclease-free water. RNA quantity and quality were measured by a NanoDrop® ND-1000 UV-Vis Spectrophotometer (Thermo Scientific, Waltham, Massachusetts) and RNAs were stored at −80 [C until use.

### cDNA synthesis and qPCR

Total RNAs were converted to cDNA by using RevertAid first strand cDNA synthesis kit (Thermo Fisher Scientific, United States) according to the manufacturer’s instructions and diluted to 5 ng/µl equivalent of total RNAs with nuclease-free water for qPCR analysis. qPCR reactions were set up as follows: 6.25 µl of GoTaq® qPCR Master Mix (Promega, Madison, WI, USA), 4.25 µl of nuclease-free water, 1 µl of 5 µM corresponding primer and 1 µl of cDNA. qPCR reactions were incubated at 95 °C for 2 min as initial denaturation, 45 cycles of denaturation at 95 °C for 15 s and annealing at 60 °C for 1 min following a melting step in a Rotor-Gene Q machine (Qiagen, Hilden, Germany). GAPDH was used for normalization. Primers were listed in Table 1.

**Table 1.**
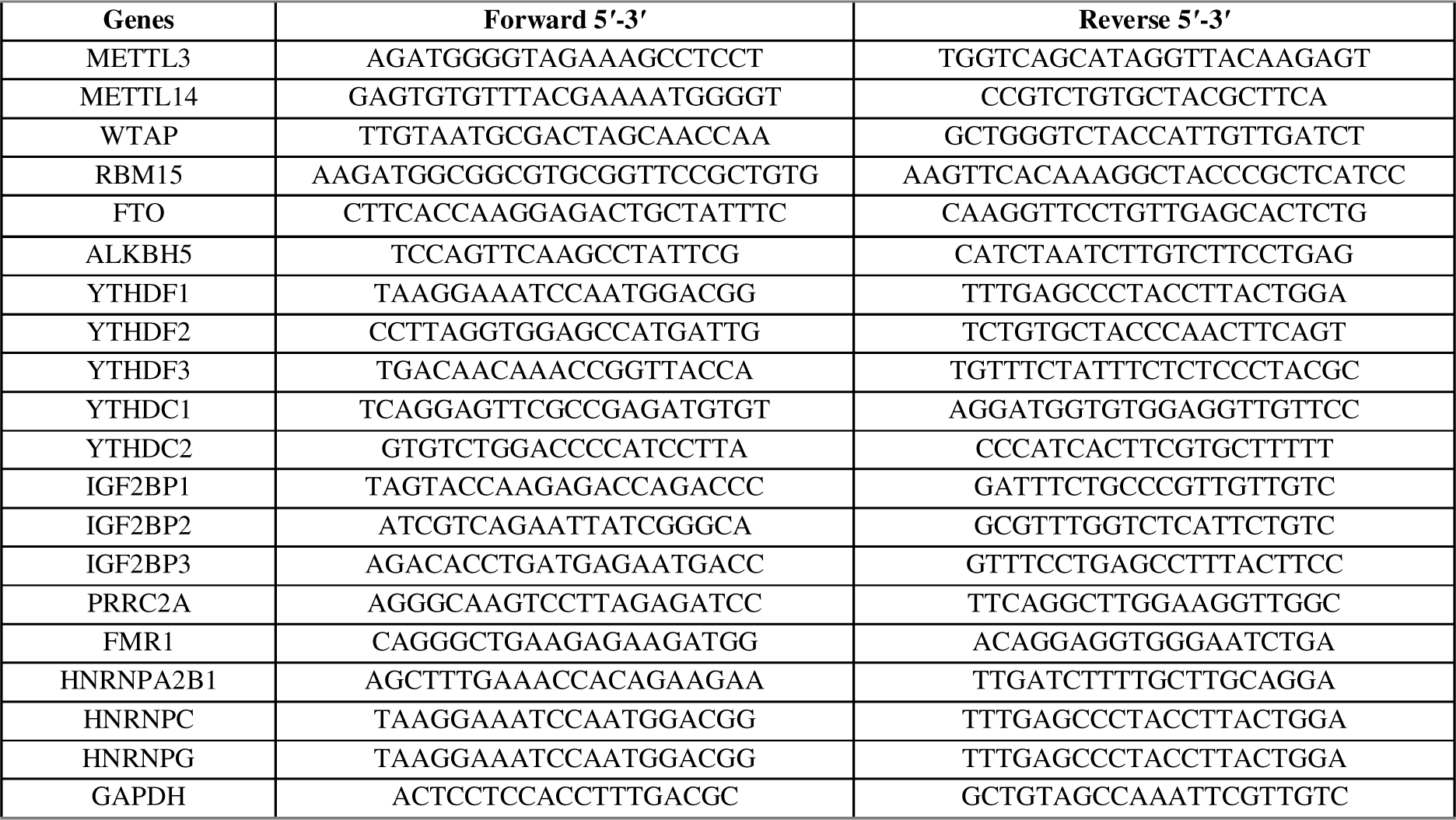
List of primers used in qPCR analysis.

### RNA-seq Data Analysis

RNA-seq data obtained from the total RNAs of cisplatin-treated HeLa cells have been published previously (Gurer et al., 2021). RNA-seq data from doxorubicin-, TNF-α and FAS ligand-treated HeLa cells are from unpublished data.

## Results

### Abundance of Transcripts of m^6^A Writers and Erasers is Deregulated in Cancer Cells

The extent of m^6^A methylation is primarily determined by a dynamic action of writers and erasers (W. Li et al., 2023). The key components of the writer complex are METTL3, METTL14, WTAP and RBM15 among others while FTO and ALKBH5 orchestrate demethylation (Luo et al., 2023). Thus, we first examined the expression levels of METTL3, METTL14, WTAP, RBM15, FTO and ALKBH5 transcripts in tumor samples and corresponding normal samples by using the GEPIA database (http://gepia.cancer-pku.cn/) (Figure 1A). Our analyses showed that the abundance of METTL3 and FTO transcripts was lower in cervical squamous cell carcinoma and endocervical adenocarcinoma (CESC), breast invasive carcinoma (BRCA), lung adenocarcinoma (LUAD) and colon adenocarcinoma (COAD) compared to their matched normal tissues. WTAP and ALKBH5 abundance displayed differences among cancer types analyzed (Figure 1A). We further examined by qPCR the abundance of m^6^A regulators in total RNAs isolated from various cell lines, namely healthy, non-metastatic and metastatic cells of cervical, breast, lung, and colon cancers. Specifically, we found lower expression of METTL3, METTL14 and WTAP transcripts while RBM15 and ALKBH5 abundance was elevated in cervical cancer cells (Figure 1B). Interestingly, RBM15 was upregulated by 20-(p<0.001) and 10-fold (p<0.01) in non-metastatic and metastatic breast cancer cells, respectively as opposed to indifferent expression based on the data in the TCGA database (Figure 1A and C). Additionally, ALKBH5 was upregulated by approximately 2.5-fold (p<0.01) in breast cancer cells (Figure 1C). In lung cancer cell lines, all m^6^A regulators analyzed were downregulated to a different extent. In particular, the transcript abundance of RBM15 and ALKBH5 were reduced by 21-fold (p<0.0001) and 4.7-fold (p<0.001), respectively, in A549 cells (Figure 1D). The abundance of RBM15 and ALKBH5 in lung cancer cell lines was negatively correlated with those in breast cancer cell lines. In colon cancer cell lines, METTL3 and RBM15 were downregulated whereas the transcript levels of FTO and ALKBH5 were upregulated (Figure 1E). We observed an increase in the RNA levels of WTAP by 8.6-fold (p<0.01) in non-metastatic cell CaCO2 whereas WTAP was downregulated by 5.2-fold (p<0.0001) in metastatic colon cancer cell line HCT116 (Figure 1E).

**Figure 1.**
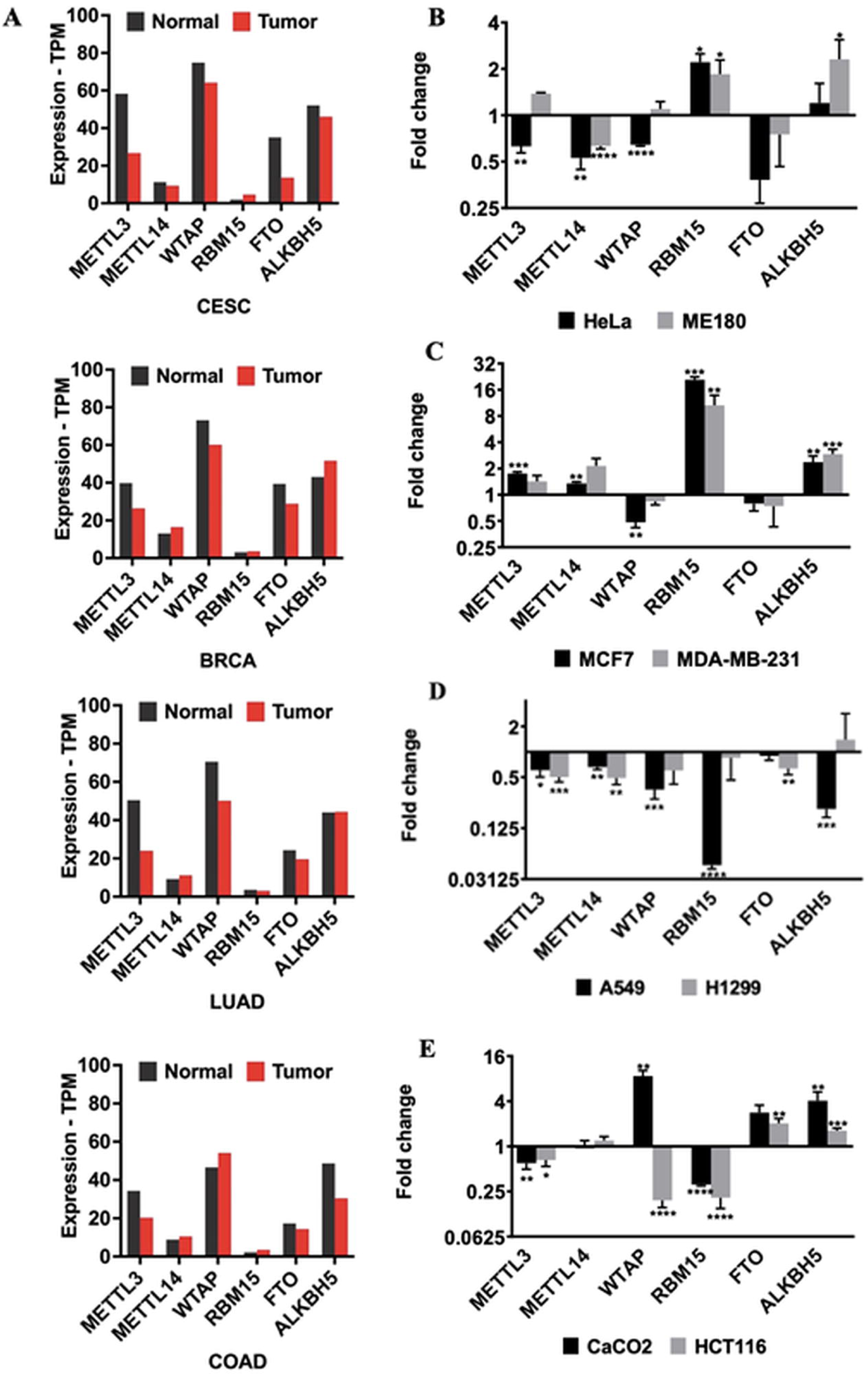
**(A)** The gene expression profile of m^6^A regulators across all tumor samples and paired normal tissues in CESC, BRCA, LUAD and COAD. The expression analysis was performed using the GEPIA2 database. CESC: Cervical squamous cell carcinoma and endocervical adenocarcinoma; BRCA: Breast invasive carcinoma; LUAD: Lung adenocarcinoma; COAD: Colon adenocarcinoma. Gene expression levels of m^6^A writers (METTL3, METTL14, WTAP and RBM15) and erasers (FTO and ALKBH5) in **(B)** HeLa and ME180, **(C)** MCF7 and MDA-MB-231, **(D)** A549 and H1299 and **(E)** CaCO2 and HCT116 cancer cell lines. All cells were normalized with their healthy cell lines. All qPCR samples were normalized with GAPDH housekeeping gene. Error bars represent meanL±LSD of three replicate samples (unpaired, two-tailed t-test). (*: p≤0.05, **: p≤0.01, ***: p≤0.001, ****: p≤0.0001)

### Expression of m^6^A RNA Methylation Regulators Under Apoptotic Conditions

One of the important hallmarks of cancer is cell death and cancer cells are notorious for their evasion of apoptotic pathways (Scheel & Schäfer, 2023). Cisplatin and doxorubicin are anti-cancer chemotherapy drugs that are widely used as inducers of the intrinsic apoptotic pathway (Motlagh et al., 2023). TNF-α and FAS ligands trigger the extrinsic apoptotic pathways by binding to their cell surface receptors (Xiao et al., 2023). To gain insight into how the expression of m^6^A regulators is affected upon initiation of apoptosis in cancer cells, we analyzed RNA-seq data to examine the expression of m^6^A regulators in HeLa cells treated with cisplatin, doxorubicin, TNF-α and FAS ligands. RNA sequencing data was obtained from HeLa cells treated with 80µM CP for 16h, 0.5 μM doxorubicin for 4 h, 0.5 μg/ml anti-Fas mAb for 16 h and 75 ng/ml TNF-α for 24 h, which was sufficient to attain an apoptosis rate of 50% in HeLa cells as published previously (Yaylak et al. 2018). The cisplatin dataset has been already published (Gürer et al., 2019). The other data sets have not been published yet (Unpublished data). By using these data sets, we analyzed the expression of a total of 18 m^6^A regulators (5 writers, 13 readers, and 2 erasers) under apoptotic conditions induced by cisplatin, doxorubicin, TNF-α and FAS ligands. Interestingly, the expression levels of regulators displayed a different pattern under all drug/ligand treatment conditions tested (Figure 2). Encouraged by the differential effect of the intrinsic and extrinsic inducers of apoptosis, we exploited cisplatin as an inducer of the intrinsic pathway and TNF-α as an inducer of the extrinsic pathway to track the expression patterns of the regulators of m^6^A RNA methylation in different cancer cell lines.

**Figure 2.**
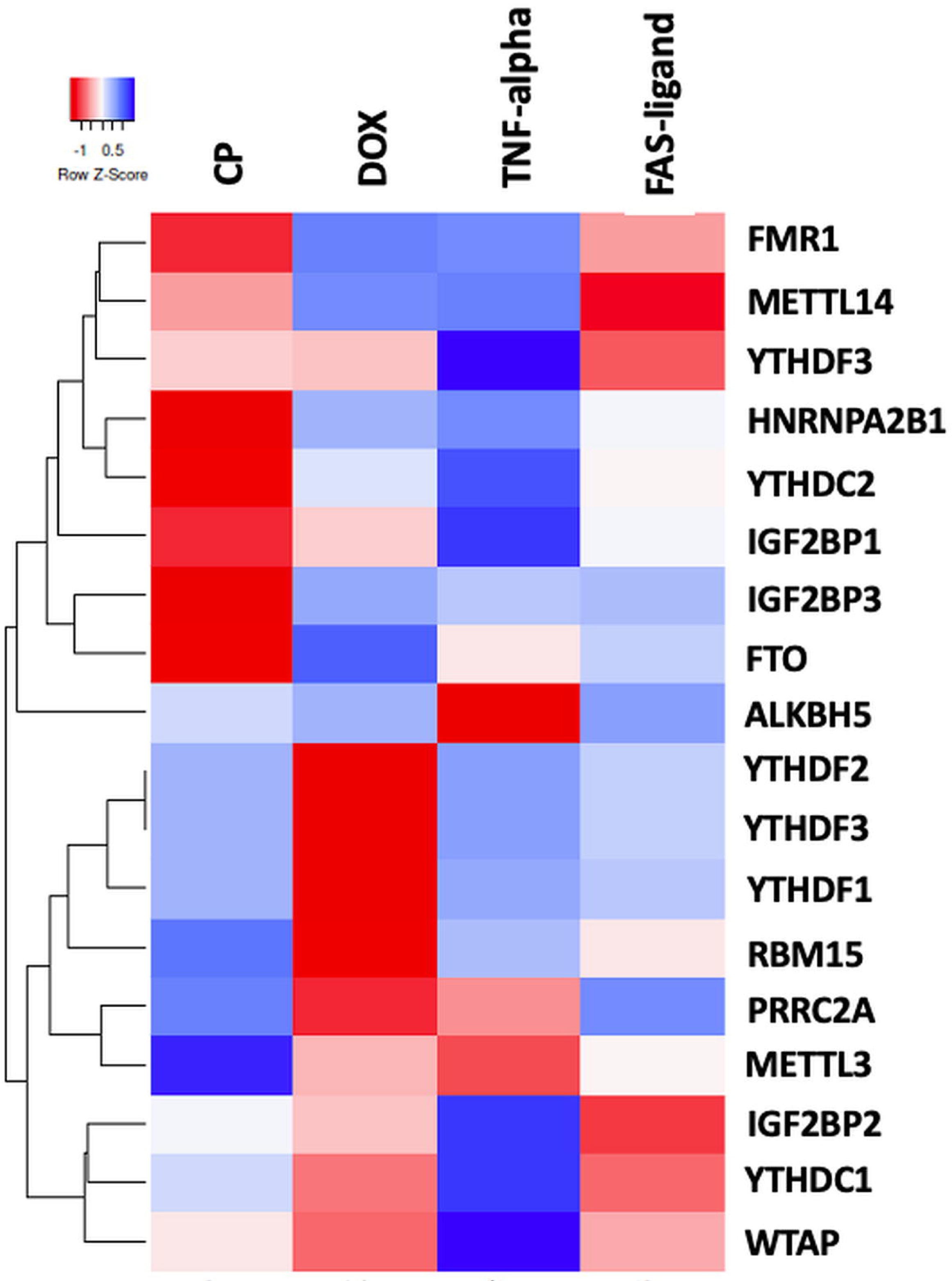
Heatmap of differentially expressed m^6^A modifiers upon the induction of the extrinsic and intrinsic pathway of apoptosis. Heatmap of RNA-seq analysis for m6A related writer, eraser and reader expressions after 4 drug treatments as CP, DOX, TNF-α and Fas ligand. Annotations on the left part of the heatmap show clustering of the m^6^A methylation related expressions.

### Expression of m^6^A regulators is perturbed by cisplatin in different cancer cell lines

In order to uncover the expression pattern of m^6^A RNA methylation regulators under cisplatin-induced apoptotic conditions, we treated HeLa, MCF7 and A549 cells with varying concentrations of cisplatin. We previously reported that 80 μM cisplatin is sufficient to induce approximately 50% of early apoptosis in HeLa cells (Yaylak et al., 2019; Figure 3A). Cisplatin at a concentration of 100 μM induced 22.8% of early apoptosis as determined by Annexin V-positive early apoptotic cells in MCF7 cells (Figure 3A). A549 cells were subjected to 80 μM cisplatin for 24 h, which was sufficient to attain an early apoptotic rate of approximately 35% compared to control DMSO (Figure 3A). We demonstrated previously that cisplatin treatment reduced the mRNA levels of METTL14 and FTO in HeLa cells by 3.3- and 6.6-fold, respectively (Alasar et al. 2022). We performed qPCR analyses with total RNAs isolated from CP-treated MCF7 and A549 cells to examine if there is a similar expression pattern in breast and lung cancer cells. Similarly, cisplatin treatment led to a significant reduction in the transcript abundance of METTL14 by 1.5-fold (p<0.05) and 1.7-fold (p<0.05) in MCF7 and A549 cells, respectively. FTO was downregulated 9.2-fold (p<0.0001) in CP-treated MCF7 and 7.7-fold (p<0.01) in CP-treated A549 cells. Also, RBM15 expression was downregulated by 1.4-fold (p<0.05) in CP-treated A549 cells (Figure 3B). We then examined the transcript abundance of m^6^A readers in HeLa, MCF7 and A549 cells upon CP treatment. Interestingly, readers displayed quite a different expression pattern although we have observed similar expression levels of writers and erasers in all cells. In CP-treated HeLa cells, YTHDF1 and YTHDF3 were upregulated by 2.6-fold (p<0.01) and 1.3-fold(p<0.05), respectively, while YTHDC2, IGF2BP2, IGF2BP3 and HNRNPA2B1 were downregulated by 2.1-fold (p<0.01), 1.3-fold(p<0.0001), 2.9-fold (p<0.01) and 1.5-fold(p<0.05), respectively (Figure 3C). In MCF7 cells, CP treatment led to a 1.7-fold (p<0.05), 7.1-fold(p<0.0001), 1.4-fold (p<0.05) and 1.9-fold (p<0.05) reduction in transcript levels of IGF2BP2, IGF2BP3, HNRNPA2B1 and HNRNPG, respectively (Figure 3C). Expression of YTHDC2, IGF2BP1, IGF2BP2, IGF2BP3, HNRNPA2B1 and HNRNPC were downregulated by 2.3-fold (p<0.05), 2.2-fold (p<0.05), 3.3-fold (p<0.001), 4.7-fold (p<0.01), 2.3-fold (p<0.05) and 1.9-fold (p<0.001), respectively, in CP-treated A549 cells (Figure 3C). All of these analyses show that cisplatin treatment leads to a reduction in the abundance of most m^6^A RNA modification regulators under our experimental conditions.

**Figure 3.**
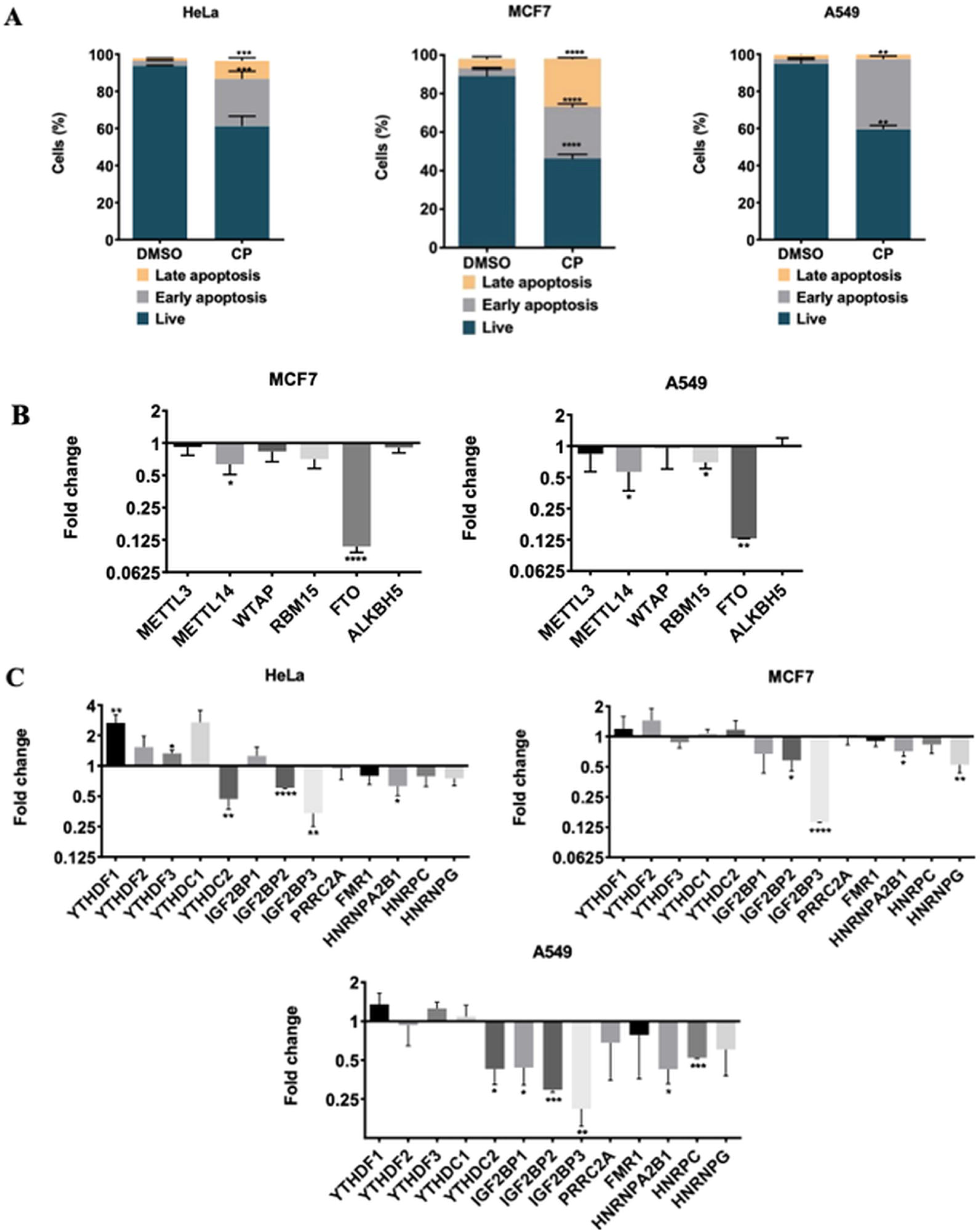
Gene expression analysis in cisplatin induced apoptotic HeLa, MCF7 and A549 cells. **(A)** HeLa, MCF7 and A549 cells were treated with 80 µM, 100 µM and 80 µM of cisplatin for 16 h, respectively. 0.1% (v/v) DMSO was used as a control. **(B)** Gene expression levels of m^6^A writers (METTL3, METTL14, WTAP and RBM15) and eraser genes (FTO and ALKBH5) in MCF7 (left) and A549 cells (right) **(C)** Gene expression levels of m^6^A readers in HeLa, MCF7 and A549 cells in CP treatment. Results were normalized against GAPDH. Experiments were conducted in triplicates. Data are presented as mean values ± SD. *: *p* ≤ 0.05, **: p≤0.01, ***: *p* ≤ 0.001, ****: *p* ≤ 0.0001 by a two-tailed unpaired t-test.

### Expressions of m^6^A regulators in TNF-***α***-treated cancer cells

Although cisplatin induces apoptosis through the intrinsic pathway, TNF-α triggers apoptosis primarily by activating the extrinsic pathway (Akçaöz-Alasar et al., 2023). We hypothesized that the abundance of m^6^A RNA regulators should be pathway-specific. To address the effects of TNF-α treatment in the abundance of m^6^A RNA regulators, we treated HeLa, MCF7 and A549 cells with TNF-α at concentration of 75 ng/ml, 10 ng/ml and 20 ng/ml with CHX of 10 µg/ml, 5 µg/ml and 10 µg/ml, respectively. TNF-α lowered the rate of live cells to 60% while causing an apoptosis rate of 23.6% in HeLa cells (Figure 4A). TNF-α induced early apoptosis rates of 14% and 33% in MCF7 and A549 cells, respectively (Figure 4A). We reported previously that, of all readers tested, merely the abundance of WTAP transcript was upregulated by 2.8-fold upon TNF-α treatment of HeLa cells (Alasar et al. 2023). We further explored the abundance of m^6^A regulators in TNF-α treated MCF7 and A549 cells. However, we have not observed any dramatic difference in the expression of m^6^A regulators, except for WTAP (1.6-fold, p<0.01) in MCF7 cells. The effect of TNF-α on the abundance of METTL3 and METTL14 transcripts was quite marginal with 1.2-fold elevation (p<0.001) (Figure 4B). It appears that METTL14 and FTO downregulations are specific to CP-induced apoptosis (Figure 3B) not TNF-α (Figure 4B). Moreover, we performed qPCR analyses to investigate the transcript level of readers under TNF-α-induced apoptotic conditions. We did not detect any discernible change in the amount of YTHDC1, YTHDC2, PRR2CA, FMR1, HNRNPA2B1 and HNRPNG in TNF-α-treated HeLa cells (Figure 4C). TNF-α treatment of MCF7 cells led to a 2.4-fold (p<0.01) and 3-fold (p<0.05) increase in the IGF2BP1 and IGF2BP3 transcript levels, respectively (Figure 4C). There was no apparent difference in the levels of readers in A549 cells (Figure 4C). Collectively, our findings showed that the expression of m^6^A regulators tend to decrease under CP-induced apoptotic conditions whereas TNF-α treatment promotes upregulation.

**Figure 4.**
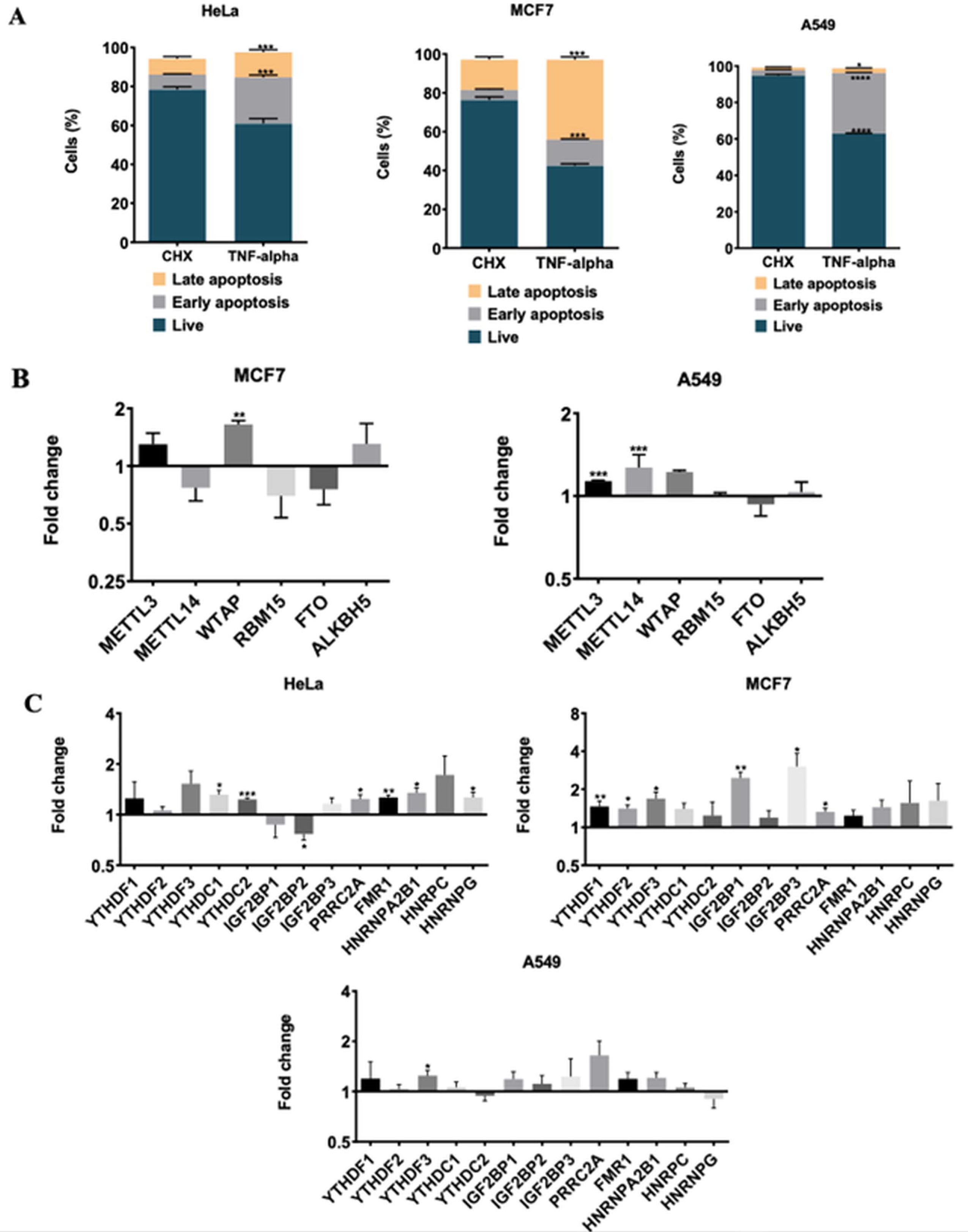
Gene expression analysis in TNF-α-induced apoptotic HeLa, MCF7 and A549 cells. **(A)** HeLa, MCF7 and A549 cells were treated with 75 ng/ml TNF-α in 5 μg/ml CHX, 10 ng/ml TNF-α in 5 μg/ml CHX and 20 ng/ml TNF-α in 10 μg/ml CHX for 24 h, respectively. CHX was used as a control. **(B)** Gene expression levels of m^6^A writers and erasers in TNF-α induced apoptotic MCF7 (left) and A549 (right) cells. **(C)** Gene expression levels of m^6^A readers in HeLa, MCF7 and A549 cells under TNF-α treated condition. All qPCR samples were normalized with GAPDH housekeeping gene. Two-tailed Student’s t test was performed to determine the statistical significance among groups. n =3 biological replicates. Data presented as mean ± SD, *: p≤0.05, **: p≤0.01, ***: p≤0.001****: p≤0.0001.

## Discussion

The most common cancer types are breast, colorectal, lung and cervix cancers according to the World Health Organization. With a diverse array of impact on tumorigenesis, m^6^A RNA methylation has recently attracted attention as an important contributor to this process. In this study, we provide expression profiles of m^6^A methylation machinery in breast, colorectal, lung and cervix cancer cell lines. CESC, BRCA, LUAD and COAD cell lines express lower amounts of METTL3 and FTO transcripts compared to their healthy counterparts, while WTAP and ALKBH5 expression differ by cancer type (Figure 1). Our analysis of cisplatin-mediated apoptosis uncovers significant reduction of METTL14 and FTO in all cell types (Figure 3). METTL14 has been reported to play a role as an oncogene in pancreatic cancer as METTL14 depletion leads to an elevated rate of apoptosis by increasing susceptibility to cisplatin in PANC-1 and CFPAC-1 cells (Kong et al.2020). METT14 may be an intermediate component in cisplatin-mediated apoptosis due to its reduction following cisplatin treatment as shown in Figure 3. Additionally, there are reports on the protective role of FTO on cisplatin-induced cytotoxicity. For example, cisplatin treatment decreases FTO expression, and downregulation of FTO enhances m^6^A methylation level and sensitization of cisplatin (Wang and Yang 2022, Zhou et al. 2019). These findings suggest that differential m^6^A methylation via dysregulation of METTL14 or FTO may aggravate cisplatin-mediated apoptosis. However, further studies are required to elucidate the common target of cisplatin in cancer types.

TNF-α treatment in our study did not display any common expression difference for writer and eraser genes (Figure 4). This observation suggests that the reduction of METTL14 and FTO may be specific to the CP-mediated intrinsic pathway. Interestingly, cisplatin treatment caused dramatically reduced expression levels of specific readers, IGF2BP2 and IGF2BP3, compared to TNF-α treatment (Figure 3-4). These findings are in parallel to the previously reported studies. Of note, ncRNA-IGF2BP2 complexes orchestrate cancer pathogenesis (Huang et al. 2018, Ma et al. 2021). Moreover, IGF2BP2 promotes the growth and metastasis of cervical cancer cells (Hu et al. 2022). Upregulation of IGF2BP2 by a miRNA-lncRNA interaction has been reported to result in increased apoptosis and decreased proliferation. Thus, a miRNA-lncRNA-IGF2BP2 axis renders cervical cancer cells resistant to cisplatin (Wu et al. 2022). A similar observation has been reported in that IGF2BP2 increases cisplatin resistance in colorectal cancer (Xia et al. 2022). Our results, on the other hand, indicate that the IGF2BP2 expression is decreased specifically upon cisplatin-induced apoptosis. Accordingly, it can be suggested that cisplatin and IGF2BP2 may have a negative correlation. Additionally, the abundance of IGF2BP3 transcript was reduced upon cisplatin treatment (Figure 3). IGF2BP3 was also reported to be associated with gastric cancer progression. IGF2BP3 is highly expressed in four gastric cancer subtypes, an indication that it may promote cell growth and invasion (Zhou et al. 2017). Yang et al. (2023) reported oncogenic and poor prognostic properties of IGF2BP3 and provided evidence that downregulation of IGF2BP3 leads to enhanced apoptosis. Additionally, IGF2BP3 has been linked to cisplatin resistance in laryngeal cancer (Yang et al. 2023), providing further evidence for the potential role of IGF2BP3 in cisplatin-mediated apoptosis.

In conclusion, although more experiments are required to delineate the molecular differences between the intrinsic and extrinsic apoptotic pathways, we report in this study differences in the abundance of the m^6^A methylation machinery in these two pathways. Since expression differences in m^6^A regulators are expected to affect the genome-wide m^6^A methylation profile, it would be important to investigate the m^6^A RNA methylome under intrinsic and extrinsic apoptotic conditions. Additionally, it begs for further experiments to delineate the importance of opposing expression of METTL14, FTO, IGF2BP2 and IGF2BP3 transcripts in cisplatin- and TNF-α-mediated apoptotic pathways.

## Acknowledgements

The authors would like to thank Özgür Okur and Murat Delman for flow cytometry analyses, and Biotechnology and Bioengineering Application and Research Centre (IZTECH, Turkey) for the instrumental help. This study is funded by the Scientific and Technological Research Council of Turkey (TUBITAK Project No: 217Z234 to BA).

## Author Contributions

BA contemplated the project. AAA, BS and İEV performed all experiments. AAA and BA wrote the manuscript, all authors read and approved the manuscript.

## Conflict of Interest

The authors declare that they have no conflict of interest.

## Notes

### Competing Interest Statement

The authors have declared no competing interest.

